# Inhibition of HIF-1α accumulation in prostate cancer cells is initiated during early stages of mammalian orthoreovirus infection

**DOI:** 10.1101/2020.09.02.279810

**Authors:** Luke D. Bussiere, Cathy L. Miller

**Affiliations:** Department of Veterinary Microbiology and Preventive Medicine, College of Veterinary Medicine, Iowa State University, Ames, Iowa 50011, USA

**Keywords:** Mammalian orthoreovirus, HIF-1α, hypoxia, prostate cancer, UV-inactivation

## Abstract

Mammalian orthoreovirus (MRV) is a safe and effective cancer killing virus that has completed Phase I-III clinical trials against numerous cancer types. While many patients experience benefit from MRV therapy, pre-defined set points necessary for FDA approval have not been reached. Therefore, additional research into MRV biology and the effect of viral therapy on different tumor genetic subtypes and microenvironments is necessary to identify tumors most amenable to MRV virotherapy. In this work we analyzed the stage of viral infection necessary to inhibit HIF-1α, an aggressive cancer activator induced by hypoxia. We ruled out a number of viral proteins and the virus genome as being necessary and determined that a step parallel with virus core movement across the endosomal membrane was required for this inhibition. Altogether, this work clarifies the mechanisms of MRV-induced HIF-1α inhibition and provides biological relevance for using MRV to inhibit the devastating effects of tumor hypoxia.

## INTRODUCTION

Viruses are generally considered disease causing agents that scientists work diligently to eliminate by vaccination; however, evidence is mounting that a select number of viruses can be exploited to improve human lives. Cancer killing, or oncolytic, viruses are wildtype (wt) or genetically engineered viruses that can be used therapeutically to locate, infect, and kill tumor cells while leaving the host unharmed (Russell et al., 2012). These oncolytic viruses have been shown to be effective in killing many tumor types in humans and other mammals by direct lysis of the cancerous cells and have also been shown to be excellent activators of antitumor immunity (Chaurasiya et al., 2018; Ungerechts et al., 2016). One such oncolytic virus is mammalian orthoreovirus (MRV) which has been shown to be safe in Phase I clinical trials and has progressed to Phase II and III clinical trials against multiple tumor types (Gong et al., 2016). While MRV is a promising cancer therapeutic, it has become clear in clinical trials that not all tumors decrease in size or dissipate following MRV therapy (Phillips et al., 2018). Therefore, to improve the efficacy of MRV as a cancer therapeutic, additional pre-clinical and clinical studies are necessary to better understand 1) the basic biology of the virus, 2) the precise effects of the virus on different tumor cell types, and 3) how different tumor environments are impacted by MRV therapy.

Tumor hypoxia is a common microenvironment in solid tumors that results in the accumulation of the alpha subunit of hypoxia-inducible factor 1 (HIF-1α) leading to poorer prognosis in cancer patients (He et al., 2016). HIF-1α is a master transcriptional regulator of cells growing in hypoxic conditions and modulates genes involved in cell growth, survival, angiogenesis, and metastasis (Semenza, 2001, 2010). In non-tumor cells many of these genes are necessary for regulation of normal growth processes, but within tumors, HIF-1α upregulation of downstream genes results in increasingly aggressive cancer growth (Vaupel, 2004). Furthermore, HIF-1α and tumor hypoxia have been shown to inhibit the immune response as well as various approved cancer drugs (Noman et al., 2015; Patiar and Harris, 2006). As a result of its critical impact on tumor progression, therapeutics that target HIF-1α are needed. Indeed, researchers have shown several benefits of HIF-1α inhibition in tumors, including resensitizing resistant tumors to chemotherapeutics and rescuing natural killer cell lysis susceptibility (Bosseler et al., 2018; Li et al., 2019).

Previously, we and others have shown that MRV inhibits the accumulation of HIF-1α *in vitro* and in tumors *in vivo* (Cho et al., 2010; Gupta-Saraf and Miller, 2014; Hotani et al., 2019; Hotani et al., 2015). MRV inhibition of HIF-1α is independent of the oxygen-dependent VHL pathway, and instead utilizes RACK1-mediated proteasomal degradation and translational inhibition to prevent HIF-1α accumulation in infected cells (Cho et al., 2010; Gupta-Saraf and Miller, 2014). Moreover, it has been demonstrated that both UV-inactivated MRV and direct introduction of MRV double-stranded RNA (dsRNA) segments into H1299 and A594 cells was sufficient to inhibit HIF-1α accumulation, in a manner independent of dsRNA recognition by the RIG-I/IPS1 pathway (Hotani et al., 2015). Apart from this, little is known about the mechanism of MRV induced inhibition of HIF-1α prompting further research into the stage of viral infection necessary for inhibition of HIF-1α to determine the extent by which MRV is useful clinically to alter the hypoxic response.

MRV infection begins when the virus attachment protein σ1 binds to JAM-A and/or sialic acid on the cell surface and is endocytosed into the cell (Barton et al., 2001; Paul et al., 1989; van den Wollenberg et al., 2012). The endosome acidifies as it progresses into a late endosome, a process that is required for efficient MRV infection (Mainou and Dermody, 2012; Sturzenbecker et al., 1987). Upon endosomal acidification, pH-dependent proteases cleave the outer capsid protein σ3, allowing further cleavage and release of μ1 fragments (Baer and Dermody, 1997; Nibert and Fields, 1992; Nibert et al., 1991). The myristoylated μ1N and ϕ fragments of μ1 penetrate the endosomal membrane forming pores that appear to be too small for viral core escape (Agosto et al., 2006; Danthi et al., 2008; Nibert and Fields, 1992; Nibert et al., 1991). It has been suggested that the sheer number of holes produced by μ1N and ϕ, or differences in osmotic pressure between the cytosol and the endosome, lead to endosomal disintegration and viral core escape (Agosto et al., 2006). However, a recent study suggests that within 4 h following MRV infection the endosome is disrupted to allow protein movement across the membrane, but the endosome remains intact (Kounatidis et al., 2020). This may indicate selective shuttling of the viral core through large pores within the endosomal membrane that shrink to preserve the integrity of the endosome (Kounatidis et al., 2020). In either case, the virus core particle escapes and resides in the cytoplasm where transcription, translation, assortment, assembly, and replication occur at or near virus formed inclusions termed viral factories (Broering et al., 2004; Dales, 1965; Desmet et al., 2014; Miller et al., 2010; Silverstein and Dales, 1968). Intact virions are then released by cell lysis or fusion of modified lysosomal compartments containing virus particles with the plasma membrane (Fernández de Castro et al., 2020).

In this work we systematically tested each step of the viral replication cycle to determine the stage at which MRV induces the inhibition of HIF-1α accumulation. Utilizing UV-inactivation and viral endosome disruption inhibitors, we identified the step during infection that was sufficient to inhibit HIF-1α, and then examined different active pathways at this stage of infection for their role in inhibition. We removed or mutated viral proteins, or dsRNA, to further investigate the mechanism and identify viral components necessary for inhibition. Finally, we compared the mechanism of UV-inactivated inhibition of HIF-1α to that of wt MRV to discern if UV-inactivated MRV mimicked what occurs during productive infection. This work has provided new insights into the mechanisms of HIF-1α inhibition by MRV and may help guide clinicians in the utilization of MRV therapy to target hypoxic tumors.

## MATERIALS AND METHODS

### Cells, viruses, antibodies, and reagents

PC3 cells were maintained in F-12K nutrient mixture Kaighn’s modification medium (Invitrogen Life Technologies) supplemented with 10% fetal bovine serum (Atlanta Biologicals), and penicillin (100 I.U./ml) streptomycin (100 μg/ml) solution (Mediatech). L929 cells were maintained in Joklik modified minimum essential medium (Sigma-Aldrich) supplemented with 2% fetal bovine serum, 2% bovine calf serum (HyClone), 2 mM L-glutamine (Mediatech), and penicillin (100 I.U./ml) streptomycin (100 μg/ml) solution. Our laboratory stocks of MRV strain type 3 Dearing Cashdollar (T3D^C^) and type 3 Dearing Fields (T3D^F^) originated from the laboratory of M. L. Nibert (Parker et al., 2002), and our stock of T3D^F^ M2 (I595K) originated from the laboratory of P. Danthi (Danthi et al., 2008). Virus was propagated and purified as previously described using Vertrel XF (DuPont) instead of Freon (Furlong et al., 1988; Mendez et al., 2000). Primary antibodies used were as follows: monoclonal mouse α-HIF-1α antibody (BD Biosciences; #610958), polyclonal rabbit α-β-Actin antibody (Cell Signaling; #4967), monoclonal rabbit α-α-Tubulin (11H10) antibody (Cell Signaling; #2125), polyclonal rabbit α-λ1 antibody (Kim et al., 2002), and rabbit α-T3D antisera (Wetzel et al., 1997). Secondary antibodies used are Alexa 488- and 594-conjugated donkey α-mouse or α-rabbit IgG antibodies (Invitrogen Life Technologies; #A-21202, #A-21207), and goat α-mouse or α-rabbit IgG-AP conjugate antibodies (Bio-Rad Laboratories, #1706520, #1706518). MG132 (Enzo Life Sciences) was used at a concentration of 50 μM, NH4Cl (Thermo Fisher Scientific) at a concentration of 20 mM, E64 (ApexBio Technology) at a concentration of 200 μM, Z-VAD-FMK (Enzo Life Sciences) at a concentration of 20 μM, and docetaxel (Acros Organics) at a concentration of 3 μM.

### Hypoxia

Cells were exposed to hypoxic conditions in a Galaxy 48R CO_2_ Incubator (New Brunswick Scientific) equipped with 1-19% O_2_ control set at 1% O_2_, 5% CO_2_, and 37°C. PC3 cells were exposed to hypoxia for 4 h (Ravenna et al., 2014).

### Infection

PC3 cells were infected with T3D^C^, T3D^F^, or I595K at a multiplicity of infection (MOI) = 20 based on cell infectious units (CIU) as described (Qin et al., 2011). Virus was UV-inactivated by exposing virus to 1J/cm^2^ of UV-irradiation (Qin et al., 2009). Since determining CIU is not possible for non-transcribing UV-irradiated virus we utilized the same number of viral particles, based on the calculation that one optical density unit at 260 nm = 2.1 × 10^12^ virions for each virus (Smith et al., 1969).

### Immunoblotting and Coomassie

PC3 cells or 1.6 × 10^6^ PFU of virus were lysed and harvested in 2x protein loading buffer (125 mM Tris-HCl [pH 6.8], 200 mM dithiothreitol [DTT], 4% sodium dodecyl sulfate [SDS], 0.2% bromophenol blue, 20% glycerol), and subject to sodium dodecyl sulfate-polyacrylamide gel electrophoresis (SDS-PAGE) for protein separation. For Coomassie staining, viral proteins separated by SDS-PAGE were stained (0.25% Coomassie Brilliant Blue R-250 [Bio-Rad Laboratories], 30% isopropanol, 20% methanol) and destained (40% methanol, 10% acetic acid). For immunoblot analysis, viral and cellular proteins separated by SDS-PAGE were transferred to nitrocellulose membranes by electroblotting, and then membranes were incubated in Tris-buffered saline (20 mM Tris, 137 mM NaCl [pH 7.6]) with 0.1% Tween 20 (TBST) with primary and secondary antibodies for 18 h and 4 h, respectively. Membranes were rinsed with TBST, treated with NovaLume Atto Chemiluminescent Substrate AP (Novus Biologicals) and imaged on a ChemiDoc XRS Imaging System (Bio-Rad Laboratories). Quantity One imaging software (Bio-Rad Laboratories) was utilized to examine and quantify the intensity of protein bands.

### RNA extraction, electrophoresis and reverse transcription

1.88 × 10^11^ viral particles of T3D^C^ and T3D^C^ top component, and UV-inactivated T3D^C^ and top component were subjected to TRIzol LS (Life Technologies) extraction via the manufacturer’s instructions. Briefly, viruses were homogenized in TRIzol LS, and the addition of chloroform separated each sample into protein, DNA, and RNA phases. The RNA phase was collected, and isopropanol and ethanol were added to precipitate and wash RNA. Extracted RNA was separated on 10% SDS-PAGE at a constant 20 mA for 10 h, and the gel was incubated in water with 3X GelRed (Phenix Research Products) for 1 h and imaged on a ChemiDoc XRS imaging system. I595K RNA was extracted via TRIzol LS extraction and subjected to RT-PCR using SuperScript IV (Invitrogen Life Technologies) as per the manufacturer’s instruction for sequencing.

### Immunofluorescence

At 24 h p.i., PC3 cells were fixed with 4% paraformaldehyde for 20 min, permeabilized with 0.2% Triton X-100 in phosphate-buffered saline (PBS) (137 mM NaCl, 3 mM KCl, 8 mM Na_2_HPO_4_, 1.5 mM KH_2_PO_4_, pH 7.4) for 5 min and blocked with 1% bovine serum albumin in PBS (PBSA) for 10 min with PBS washes between each step. Cells were then incubated for 45 min at room temperature with primary antibodies diluted in PBSA and washed two times with PBS, followed by incubation with secondary antibody diluted in PBSA for 45 min and two additional PBS washes. Coverslips with labeled cells were mounted with ProLong Gold antifade reagent with DAPI (4,6-diamidino-2-phenylindole dihydrochloride; Invitrogen Life Technologies) on slides. Each coverslip was then examined on a Zeiss Axiovert 200 inverted microscope equipped with fluorescence optics to determine the number of cells expressing nuclear HIF-1α. Representative pictures were taken by a Zeiss AxioCam MR color camera using AxioVision software (4.8.2). Images were prepared using Adobe Photoshop and Illustrator software (Adobe Systems).

### Apoptosis detection

Apoptosis induction by T3D^C^ and inhibition by Z-VAD-FMK was determined using the Caspase-Glo 3/7 Assay System (Promega). PC3 cells were plated at 1 × 10^4^ cells per 96 well plate and 24 h later were either treated with docetaxel, mock-infected, or infected with T3D^C^ or UVT3D^C^, and were then treated with or without Z-VAD-FMK. At 24 h p.i. media was removed and 50 μl fresh media and 50 μl Caspase-Glo Reagent was added to each well. After 30 min incubation at room temperature, the plate was read on a Glomax Multi Detection Plate Reader (Promega).

## RESULTS

### MRV gene expression is not required for inhibition of HIF-1α accumulation in PC3 cells

Previous investigation into MRV-induced inhibition of HIF-1α has suggested that UV-inactivated MRV, which is deficient in virus life cycle events downstream of and including mRNA transcription, inhibits HIF-1αto wt MRV levels in H1299 and A594 human lung cells *in vitro* (Hotani et al., 2015). We have previously shown that MRV inhibits HIF-1αin prostate cancer (PCa) cell lines (Gupta-Saraf and Miller, 2014), and reasoned UV-inactivated MRV may also inhibit HIF-1α in these cells. To determine the extent by which UV-inactivation impacts MRV inhibition of HIF-1α accumulation in PCa cells, we mock-infected or infected PC3 cells at an MOI = 20 with MRV strain T3D^C^ or UV-inactivated T3D^C^ (UVT3D^C^). At 20 h post-infection (p.i.) we exposed a subset of cells to hypoxic conditions (1% O_2_, 5% CO_2_ at 37°C) for 4 h before harvesting and subjecting to immunoblot assay. To confirm that UV-inactivation was sufficient to interfere with viral gene expression, we immunostained with antibodies against viral structural (λ1) and non-structural (μNS) proteins. As expected, λ1 was detected in UVT3D^C^ and, to a greater extent in T3D^C^ infected cells; however, only T3D^C^ expressed μNS, suggesting that UVT3D^C^ was effectively inactivated and unable to transcribe and translate protein (Fig. 1A). Immunostaining against HIF-1α revealed that both T3D^C^ and UVT3D^C^ inhibited accumulation of HIF-1α under hypoxic conditions. Quantification of immunoblot replicates demonstrated that HIF-1α accumulation was significantly less (*P*< 0.0001) in T3D^C^ and UVT3D^C^ infected cells compared to mock-infected cells (Fig. 1B). To strengthen this finding, we also performed immunofluorescence assays of cells infected with T3D^C^ and UVT3D^C^. Immunofluorescence showed that in cells infected with either T3D^C^ or UVT3D^C^, there was little to no HIF-1α accumulation (Fig. 1C-D). Interestingly, in both the immunoblot and immunofluorescence analysis, there was significantly less HIF-1α accumulation in UVT3D^C^ infected cells compared to T3D^C^ infected cells. Taken together, this data suggests that MRV infection inhibits HIF-1α accumulation in the absence of gene expression, and further, that UV-inactivated MRV has the capacity to inhibit HIF-1α accumulation in PCa cells with increased efficiency compared to wt virus.

**Figure 1.**
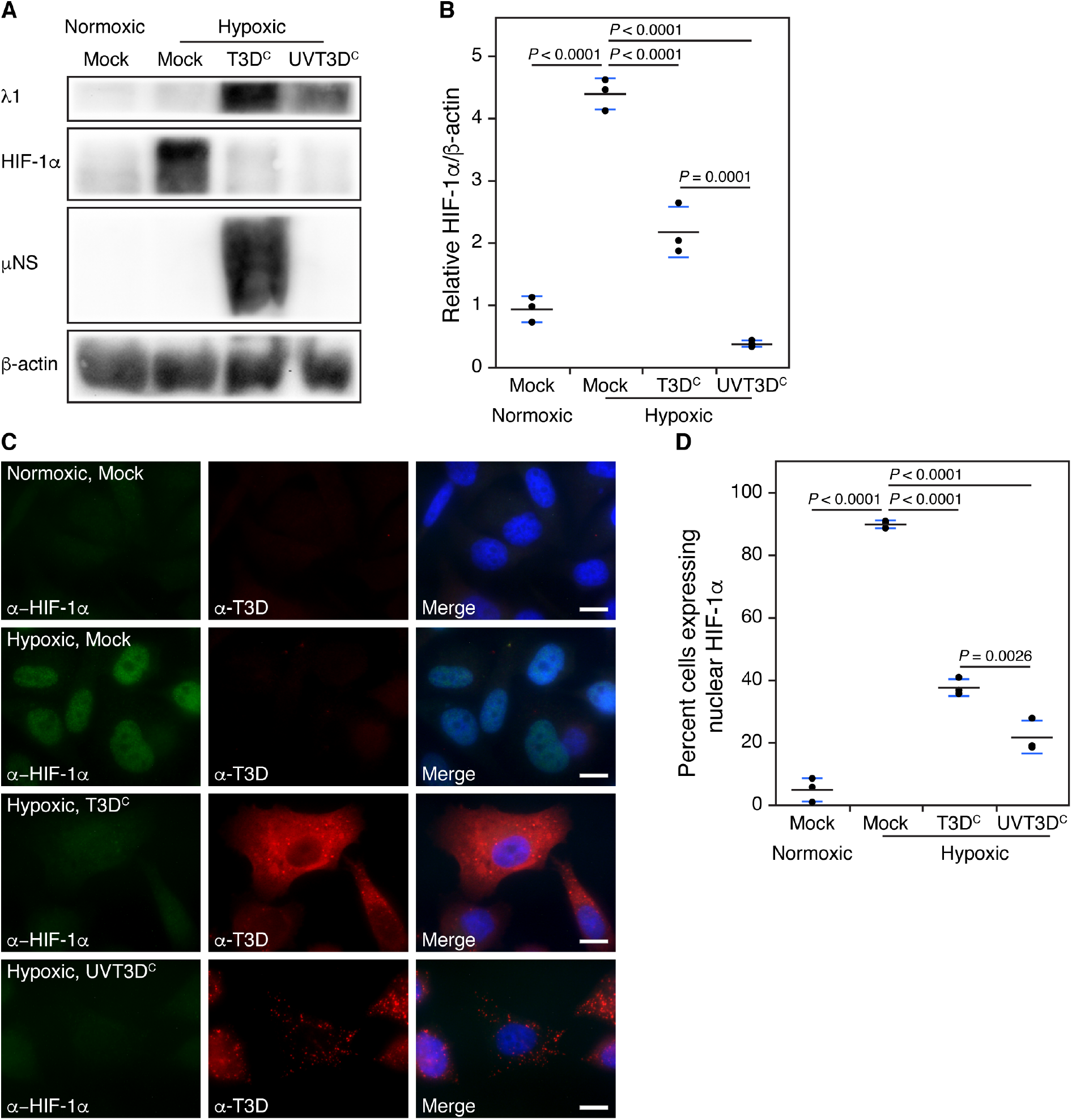
MRV gene expression is not necessary for HIF-1α inhibition. PC3 cells were mock-infected or infected at an MOI= 20 with T3D^c^ or an equal number of viral particles of T3D^c^ exposed to 1 J/cm^2^ of UV-irradiation (UVT3D^c^). At 20 h p.i. cells were exposed to normoxic or hypoxic conditions for 4 h and were collected for immunoblot or immunofluorescence analysis. (A) Cell lysates were run on SOS-PAGE, transferred to nitrocellulose and stained with antibodies against the virus structural protein λ1, the nonstructural μNS, HIF-1α and β-actin. (B) lmmunoblots were analyzed on Quantity One software to quantify HIF-1α/β-actin protein levels. (C) Cells were fixed for immunofluorescence analysis with primary α-HIF-1α antibody, α-T3D antisera, and secondary Alexa 594-conjugated donkey a-rabbit or Alexa 488-conjugated donkey α-mouse lgG. A merged image is also shown with DAPI staining. Images are representative of the observed phenotype. Bars= 10 μm. (D) Ten fields of views of cells were counted, and the number of cells expressing nuclear HIF-1α is shown. Black dots represent each replicate, the black line represents the average and the blue lines represent the standard deviation. *P* values were determined using ANOVA with Tukey’s multiple comparison test in JMP® Pro 14.2.0.

### Viral capsid cleavage, endosomal escape, or endosomal disruption is necessary for MRV-induced inhibition of HIF-1α accumulation

Our previous data suggested that viral transcription and subsequent steps are not required for HIF-1α inhibition; therefore, we next examined events directly upstream of viral transcription in the MRV life cycle, inclusive of outer capsid protein cleavage and escape of the viral core particle from the endosome. PC3 cells were subjected to NH_4_Cl or E64, which block viral capsid protein cleavage and virus core escape from the endosome by preventing pH change within the endosome necessary for activation of proteases (NH_4_Cl) or directly inhibiting cysteine proteases necessary for capsid protein cleavage (E64) (Sturzenbecker et al., 1987). At 4 h post drug addition, PC3 cells were mock-infected or infected with T3D^C^. At 20 h p.i., cells were placed under hypoxic conditions for 4 h, then harvested and subjected to immunoblot analysis (Fig. 2A). Similar to prior experiments, blots were immunostained with antibodies against μNS and λ1 to show that cells were infected (λ1) but were unable to escape the endosome to begin transcribing and translating protein (μNS). As expected, we observed significant HIF-1α inhibition in T3D^C^ infected cells compared to mock-infected cells under hypoxia; however, there was not a significant difference in accumulated HIF-1α levels in infected cells treated with NH4Cl or E64 compared to drug-treated mock-infected cells (Fig. 2B). To determine if HIF-1α inhibition mediated by UVT3D^C^ was also sensitive to NH_4_Cl and E64 treatment, we repeated the previous experiment with cells infected with UVT3D^C^ and observed similar results (Fig. 2C-D). Lastly, our results were confirmed via immunofluorescence where T3D^C^ and UVT3D^C^ infected cells inhibited HIF-1α accumulation in the absence of drug treatment and failed to inhibit HIF-1α when NH_4_Cl or E64 were added (Fig. 2E). Together our data suggests that a step between viral outer capsid cleavage in the endosome and transcription during T3D^C^ infection is sufficient to inhibit HIF-1α accumulation in PCa cells.

**Figure 2.**
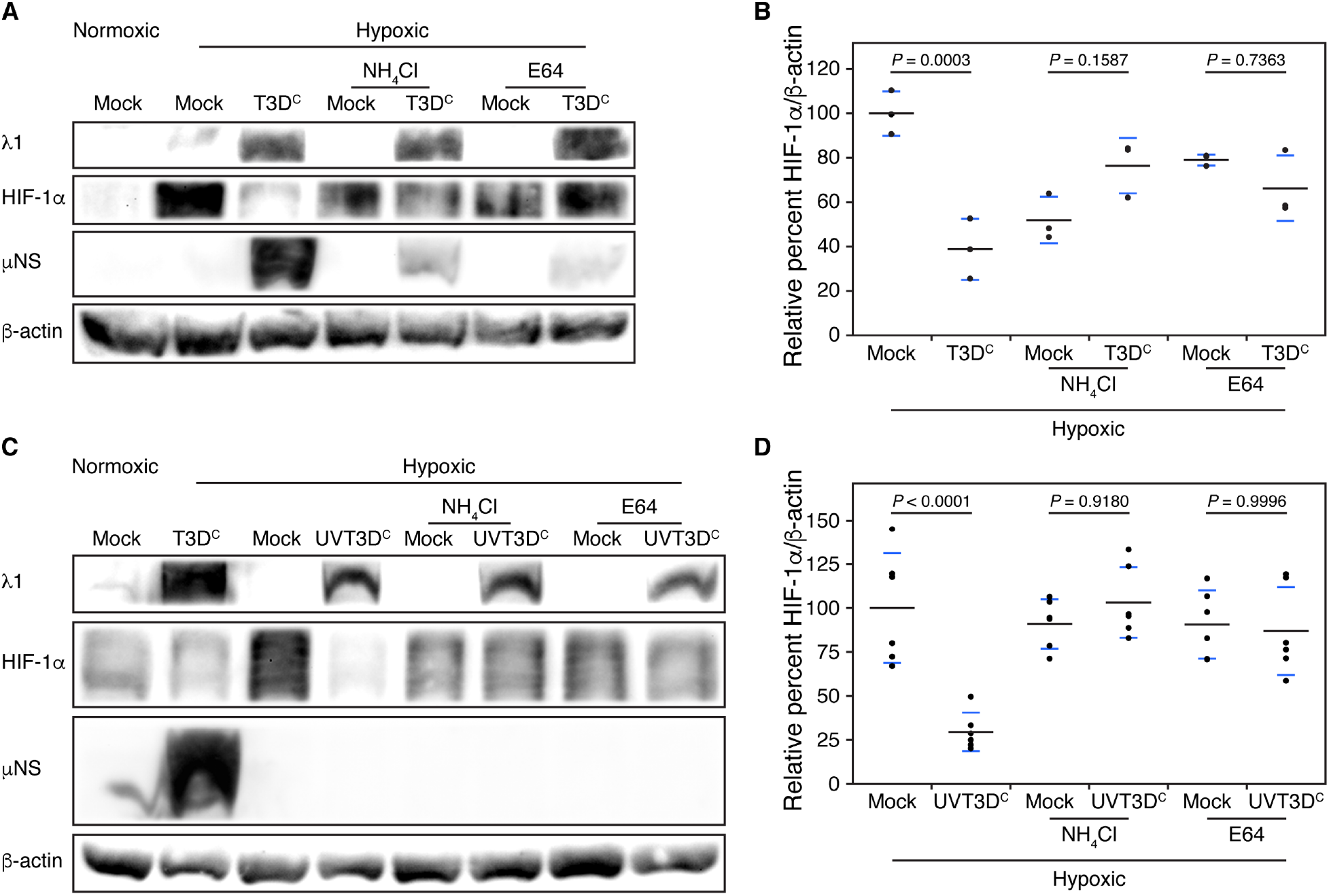
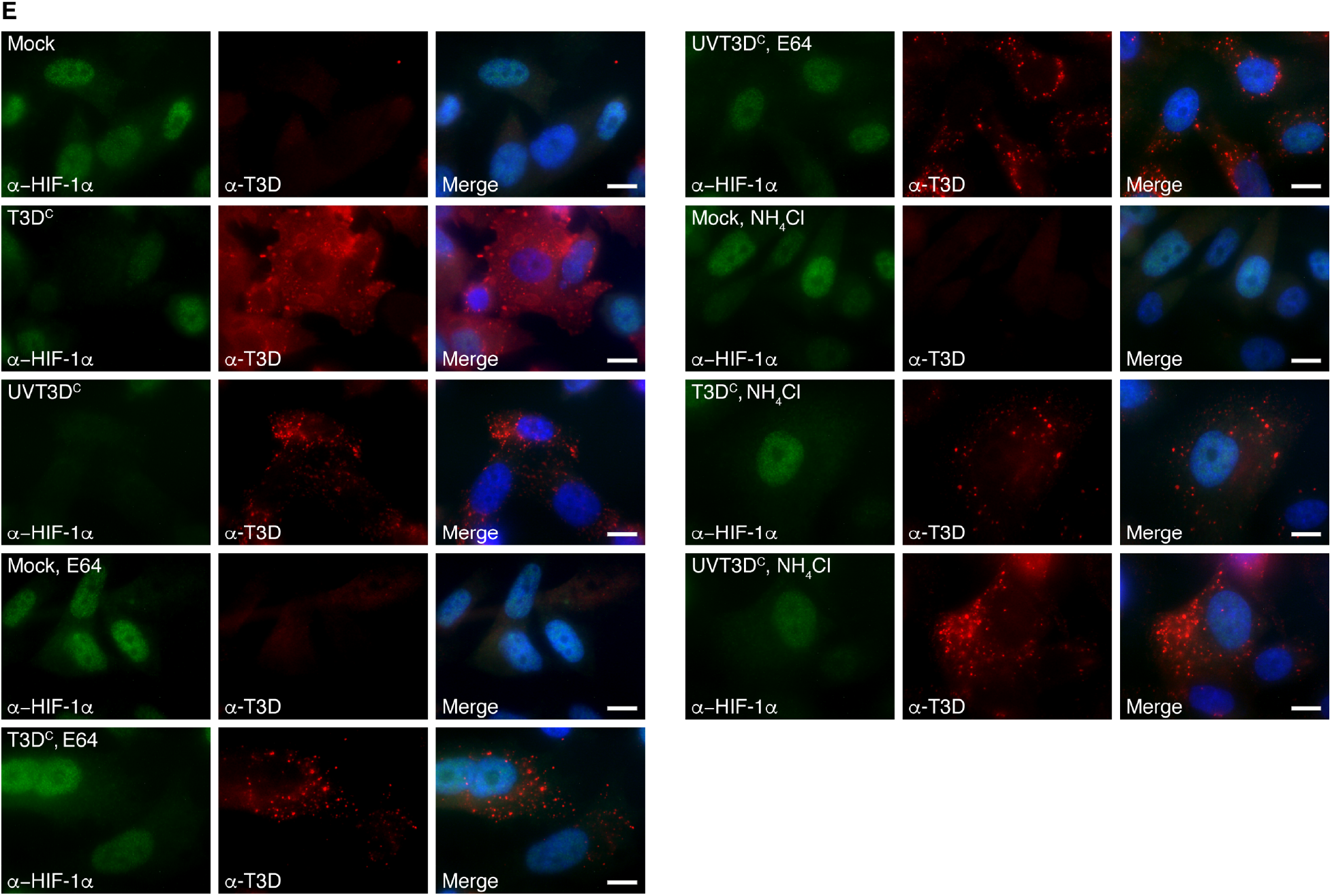
Viral outer capsid cleavage inhibitors rescue HIF-1α accumulation in MRV infected PC3 cells. PC3 cells were treated with NH_4_CI (200 mM) or E64 (20 μM) 4 h p.i. and were then mock-infected or infected at an MOI= 20 with T3D^c^ or an equal number of viral particles of T3D^c^ exposed to 1 J/cm^2^ of UV-irradiation (UVT3D^c^). At 20 h p.i. cells were exposed to normoxic or hypoxic conditions for 4 h and were collected for immunoblot or immunofluorescence analysis. Cell lysates from T3D^c^ infected PC3 cells (A) or UVT3D^c^ infected PC3 cells (C) were run on SOS-PAGE, transferred to nitrocellulose and stained with antibodies against the virus structural protein A1, the nonstructural μNS, HIF-1 α and β-actin. lmmunoblots of T3D^c^ (B) or UVT3D^c^ (D) infected cells were analyzed on Quantity One software to quantify HIF-1α/β-actin protein levels. Black dots represent each replicate, the black line represents the average and the blue lines represent the standard deviation. *P* values were determined using ANOVA with Tukey’s multiple comparison test in JMP® Pro 14.2.0. (E) Cells were fixed for immunofluorescence analysis with primary α-HIF-1α antibody, α-T3D antisera, and secondary Alexa 594-conjugated donkey α-rabbit or Alexa 488-conjugated donkey α-mouse lgG. A merged image is also shown with DAPI staining. Images are representative of the observed phenotype. Bars= 10 μm.

### Outer capsid protein σ3 is not necessary for T3D^C^-induced HIF-1α inhibition

Data supports a hypothesis where MRV core escapes the acidified late endosome following protease cleavage and removal of the σ3 protein from the virion, and membrane disruption by the newly exposed hydrophobic portions of μ1 (Agosto et al., 2006; Baer and Dermody, 1997; Danthi et al., 2008; Kounatidis et al., 2020; Mainou and Dermody, 2012; Nibert and Fields, 1992; Nibert et al., 1991; Sturzenbecker et al., 1987). Therefore, there are viral components that are released from the virion into the endosome as a result of capsid protein cleavage, and presumably from the endosome into the cytoplasm during endosomal disruption including the viral core particle, σ1, σ3 (or σ3 cleavage fragments), μ1 (or μ1 cleavage fragments), and dsRNA as a result of breakdown of the core (Sherry, 2009). Since there are 600 copies of σ3 and μ1 per virion we began investigating the role of these two proteins on HIF-1α inhibition beginning with σ3 (Dryden et al., 1993).

While the outermost capsid protein, σ3, provides stability to the viral capsid it is not required for infection and can be removed by proteases *in vitro* (Borsa et al., 1973; Drayna and Fields, 1982; Joklik, 1972; Shatkin and LaFiandra, 1972). Therefore, immediately prior to PC3 infection, T3D^C^ was incubated with α-chymotrypsin for 5 min to cleave σ3 to produce replication-competent infectious subvirion particles (ISVPs). A portion of the ISVPs was then exposed to UV light to produce UVISVPs as done previously to make UVT3D^C^. T3D^C^, UVT3D^C^, ISVP, and UVISVP samples were separated on SDS-PAGE and stained with Coomassie to confirm successful ISVP production through demonstration of σ3 degradation and cleavage of μ1 into the δ fragment (Fig. 3A) as previously shown (Nibert and Fields, 1992). PC3 cells were then infected for 20 h, followed by 4 h hypoxia treatment. Immunoblots of these samples demonstrated that both ISVPs and UVISVPs were able to significantly inhibit HIF-1α accumulation compared to mock-infected cells (Fig. 3B-C). Since UVISVPs lack σ3 and are unable to transcribe and translate σ3 but were still able to inhibit HIF-1α accumulation, this suggests that σ3 is dispensable for HIF-1α inhibition. Furthermore, immunofluorescence assays of T3D^C^, ISVP, and UVISVP infected cells confirmed that HIF-1α accumulation was inhibited in hypoxic cells infected with ISVPs and UVISVPs (Fig. 3D).

**Figure 3.**
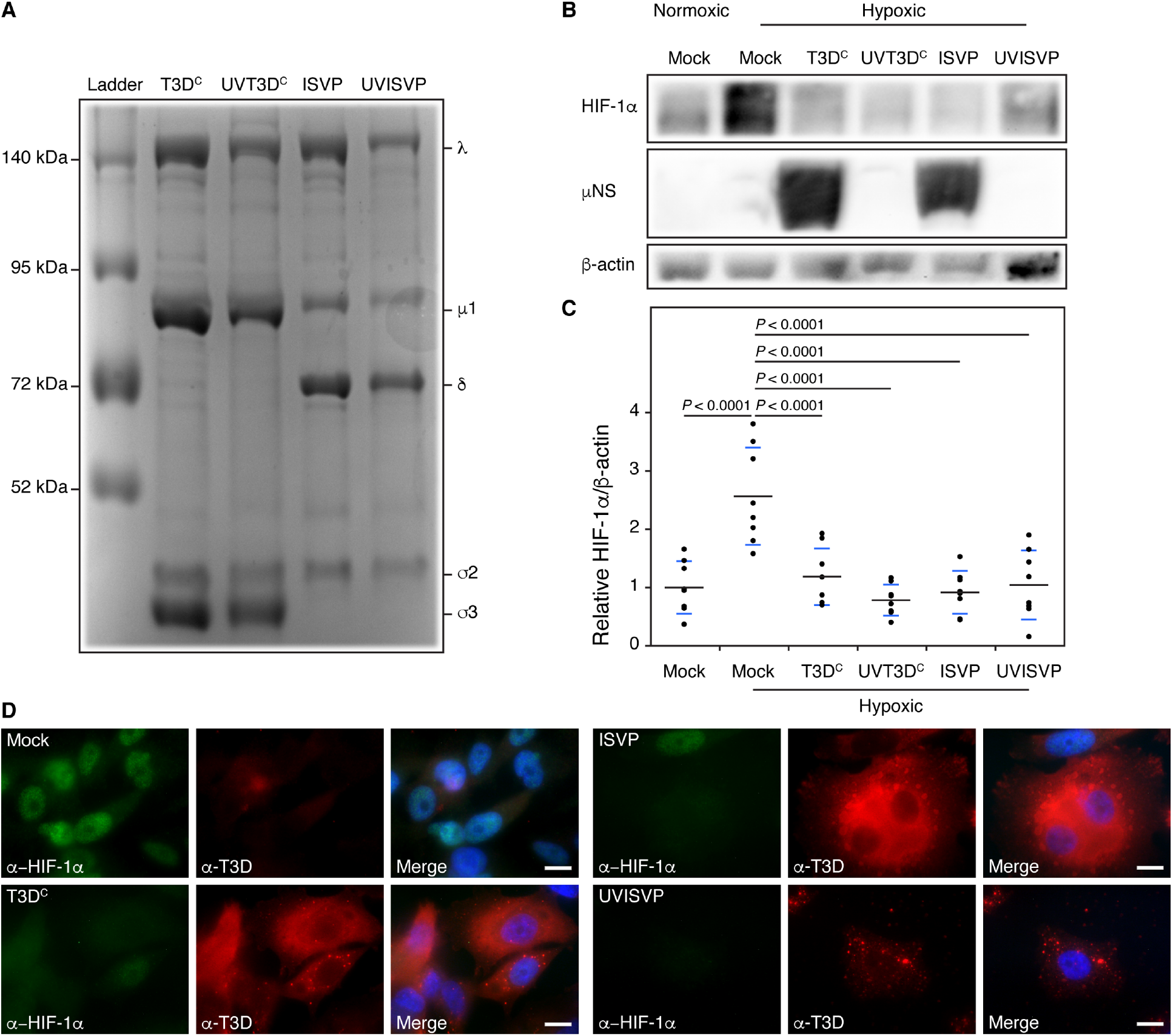
ISVPs inhibit HIF-1α similar to wt virus. 1.6 × 10^7^ Cl Us of T3D^c^ virus was treated with 200 μM α-chymotrypsin for 5 min to cleave the outer capsid proteins σ3 and μ 1 to form ISVPs. T3D^c^ and ISVP were further subjected to 1 J/cm2 UV-irradiation (UVT3Oc and UVISVP). (A) Equal viral particles of each T3D^c^, UVT3D^c^, ISVP and UVISVP were run on SOS-PAGE and stained with Coomassie to observe MRV structural proteins. ISVP production is confirmed by the cleavage of o3 and μ1 to δ as shown. PC3 cells were mock-infected or infected at an MOI = 20 with T3D^c^ and an equal number of viral particles of UVT3D^c^, ISVP or UVISVP. At 20 h p.i. cells were exposed to normoxic or hypoxic conditions for 4 h and were collected for immunoblot or immunofluorescence analysis. (B) Cell lysates were run on SOS-PAGE, transferred to nitrocellulose and stained with antibodies against the virus nonstructural protein μNS, HIF-1α, and β-actin. (C) lmmunoblots were analyzed on Quantity One software to quantify HIF-1α/β-actin protein levels. Black dots represent each replicate, the black line represents the average and the blue lines represent the standard deviation. *P* values were determined using ANOVA with Tukey’s multiple comparison test in JMP® Pro 14.2.0. (D) Cells were fixed for immunofluorescence analysis with primary α-HIF-1α antibody, α-T3O antisera, and secondary Alexa 594-conjugated donkey α-rabbit or Alexa 488-conjugated donkey α-mouse lgG. A merged image is also shown with OAPI staining. Images are representative of the observed phenotype. Bars = 10 μm.

### Apoptosis-defective MRV inhibits HIF-1α accumulation

We next turned our attention to the membrane penetration protein, μ1, which plays multiple roles during MRV infection. As a result of cleavage of σ3 by endosomal proteases, μ1 is exposed and undergoes autocatalytic cleavage resulting in release of active fragments of the protein. Both the myristoylated μ1N and the ϕ fragment poke holes in the endosomal membrane leading to core particle release into the cytoplasm (Agosto et al., 2006; Danthi et al., 2008; Nibert and Fields, 1992; Nibert et al., 1991). Furthermore, ϕ has also been shown to induce cellular apoptosis (Coffey et al., 2006). It is not possible to mutate sites involved in the membrane penetrating function of μ1N and maintain virus infectivity. However, previously described mutants in the ϕ region of μ1 (I595K) of the T3D^F^ strain of MRV have been found to decrease MRV’s capacity to induce apoptosis while maintaining wt levels of replication (Danthi et al., 2008). We took advantage of this mutant to examine the role of μ1 induction of apoptosis in HIF-1α inhibition. MRV virus strains T3D^F^ and T3D^C^ are genotypically quite similar, however, to prove that differences within the two strains do not result in differences in HIF-1α inhibition we compared HIF-1α accumulation in I595K, T3D^F^, and T3D^C^ infected cells. To determine the extent by which incoming μ1-induced apoptosis contributes to the inhibition of HIF-1α, the three strains were UV-inactivated. At 20 h p.i. cells were placed under hypoxia for 4 h, harvested, lysed, and subjected to immunoblot analysis using antibodies against λ1, HIF-1α, μNS, and β-actin (Fig. 4A). Immunoblot data was quantified and UVT3D^C^, UVT3D^F^ and UVI595K all showed significant inhibition of HIF-1α accumulation compared to mock-infected cells under hypoxia (Fig. 4B). There was no significant difference in HIF-1α accumulation between UVT3D^F^ and UVI595K infected cells, suggesting the apoptosis function of μ1, which is maintained in UV-inactivated MRV infected cells (Tyler et al., 1995), is not required for inhibition of HIF-1α accumulation during infection.

**Figure 4.**
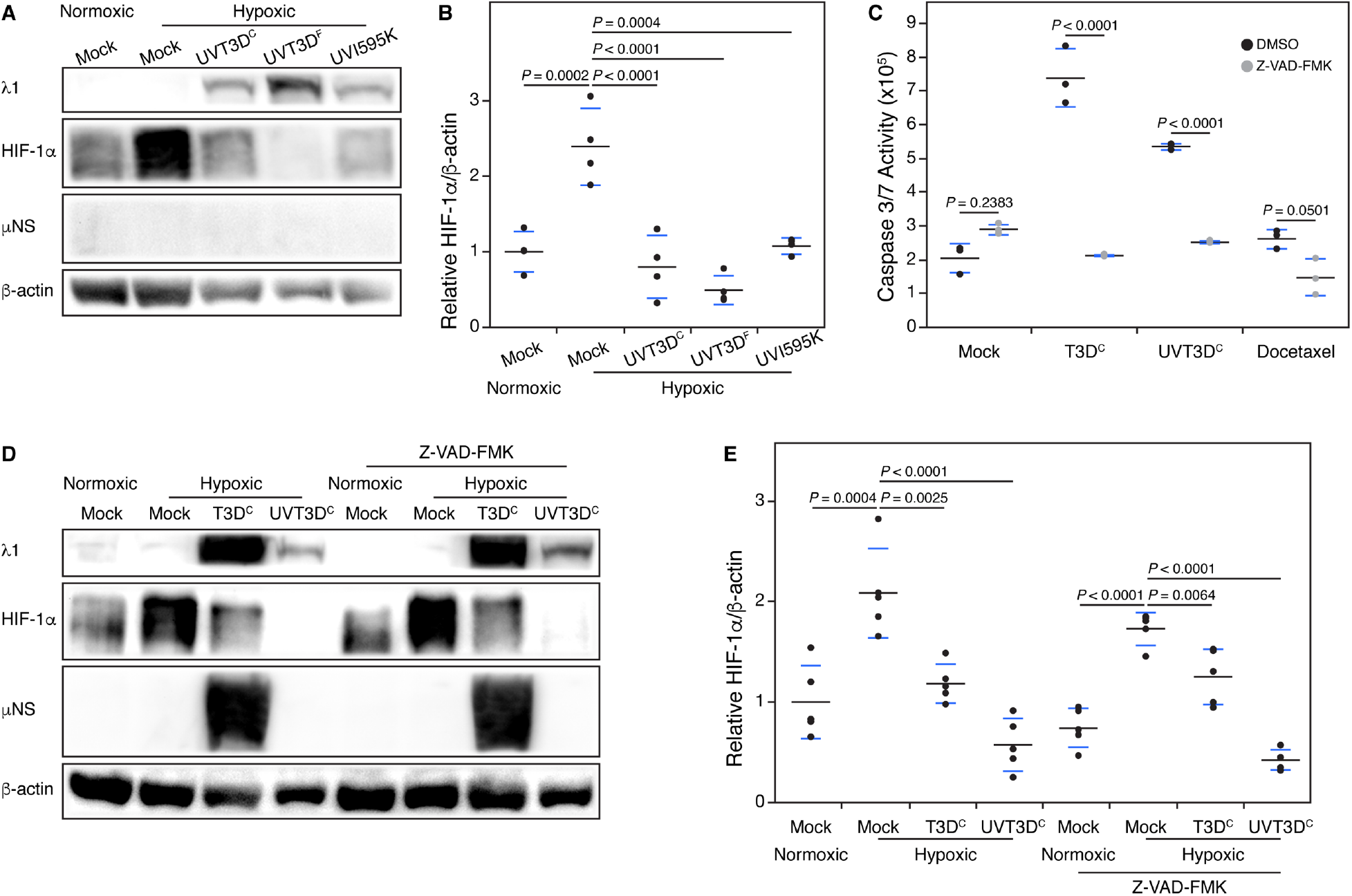
μ1 induced apoptosis does not contribute to HIF-1α inhibition. T3D^c^, T3D^F^, and I595K viruses at 1.6 × 10^7^ Cl Us were UV-irradiated at 1J/cm^2^. PC3 cells were mock-infected or infected with UVT3D^c^, UVT3D^F^, or UVl595K, and at 20 h p.i. cells were exposed to normoxic or hypoxic conditions for 4 h and were collected for immunoblot analysis. (A) Cell lysates were run on SOS-PAGE, transferred to nitrocellulose and stained with antibodies against the virus structural protein λ1, the nonstructural μNS, HIF-1α, and β-actin. **(B)** lmmunoblots were analyzed on Quantity One software to quantify HIF-1α/β-actin protein levels. PC3 cells were mock-infected or infected with T3D^c^ or UVT3D^c^, with or without treatment of the pan-caspase inhibitor Z-VA D-FMK. At 20 h p.i. and treatment cells were exposed to normoxic or hypoxic conditions for 4 h and (C) were subject to caspase 3/7 activity assay along with treatment of the apoptosis inducing docetaxel (3 μM), or (D) were collected for immunoblot analysis as described in (A). **(E)** lmmunoblots were analyzed as describe in (8). Black dots represent each replicate, the black line represents the average and the blue lines represent the standard deviation. *P* values were determined using ANOVA with Tukey’s multiple comparison test in JMP® Pro 14.2.0.

To strengthen this hypothesis, we utilized the pan-caspase inhibitor Z-VAD-FMK, which binds and inhibits caspase proteases needed for apoptosis induction, in combination with T3D^C^ to observe the impact of virus-induced apoptosis on HIF-1α accumulation. To confirm that Z-VAD-FMK at 20 μM inhibits apoptosis we mock-infected or infected PC3 cells with T3D^C^ or UVT3D^C^, or treated cells with the apoptosis inducing docetaxel, and then treated cells with or without Z-VAD-FMK. After 20 h p.i. and treatment, cells were exposed to hypoxia for 4 h and then caspase 3/7 activity was quantified (Fig. 4C). In T3D^C^, UVT3D^C^, and docetaxel infected/treated cells, caspase 3/7 activity, indicative of apoptosis induction, was increased compared to mock-infected cells. When cells were exposed to Z-VAD-FMK, caspase 3/7 activity was decreased suggesting that Z-VAD-FMK at 20 μM is sufficient to inhibit viral induced apoptosis in PC3 cells. Next, PC3 cells were mock-infected or infected, treated with or without Z-VAD-FMK, and placed under hypoxia as previously described for caspase 3/7 activity assay. Cells were harvested and lysed for immunoblot analysis, and in cells treated with Z-VAD-FMK, both T3D^C^ and UVT3D^C^ were able to inhibit HIF-1α similar to untreated cells (Fig. 4D). Quantification of immunoblot replicates demonstrated that T3D^C^ and UVT3D^C^ were able to significantly inhibit HIF-1α accumulation with or without the treatment of Z-VAD-FMK (Fig. 4E). Together these experiments suggest that μ1-or MRV-induced apoptosis does not contribute significantly to the inhibition of HIF-1α accumulation.

### Reovirus dsRNA does not significantly contribute to HIF-1α inhibition during viral infection

In addition to the outer capsid proteins, dsRNA is also a major component of entering virus particles. A previous report suggested transfected MRV dsRNA as well as polyI:C induce inhibition of HIF-1α accumulation (Hotani et al., 2015). However, naked dsRNA delivery via transfection differs significantly from dsRNA genome delivered within the protective viral core during natural infection; therefore, we devised an experimental approach that more closely recapitulates MRV infection to address the role of genomic dsRNA in HIF-1αinhibition. Upon cesium chloride (CsCl) gradient purification of MRV, a top and bottom component band form, where the bottom component consists primarily of virus containing a full complement of the ten viral genome segments, while the top band is considered genome deficient (Smith et al., 1969). For these experiments, we collected both the bottom component (T3D^C^) and the top component (TC) from the same CsCl gradient. We first extracted dsRNA from the same number of viral particles, calculated from absorbance at 260 nm (A260) = 2.1 × 10^12^ particles, of T3D^C^ and TC, and separated the dsRNA on SDS-PAGE (Smith et al., 1969). These experiments demonstrated that when the same number of particles were examined there was a substantial decrease in dsRNA within the TC compared to T3D^C^ (Fig. 5A). Next, 1.88 × 10^12^ viral particles of each T3D^C^ and TC were either left untreated or UV-treated. T3D^C^, TC, UVT3D^C^, and UVTC were lysed, separated on SDS-PAGE, and stained with Coomassie to examine virus structural proteins (Fig. 5B). The amount of each observed viral protein on Coomassie was similar between T3D^C^ and TC, and UVT3D^C^ and UVTC, suggesting that a similar number of TC and T3D^C^ virions (Fig. 5B) result in decreased dsRNA in TC compared to T3D^C^ (Fig. 5A). To determine the effect of dsRNA on HIF-1α accumulation during MRV infection, PC3 cells were infected with 1.88 × 10^12^ particles, equivalent to an MOI = 20 in T3D^C^ infected cells as previously used, of T3D^C^, TC, UVT3D^C^, and UVTC. At 20 h p.i., cells were grown under hypoxia for 4 h, then harvested, lysed, and immunoblotted with antibodies against λ1, HIF-1α, μNS, and β-actin. The immunoblot and quantified data (Fig. 5C-D) show that the dsRNA deficient TC was able to inhibit HIF-1α to similar levels to T3D^C^, suggesting dsRNA does not play a role in the inhibition of HIF-1α accumulation during normal infection. Interestingly, we observed a repeatable decrease in HIF-1α inhibition in UVTC relative to UVT3D^C^ infected cells; however, this decrease did not reach significance over five biological replicates. Altogether this suggests that dsRNA may play a minor role in inhibiting HIF-1α when the virus is unable to progress into stages of its replication cycle beyond entry, but not during active infection.

**Figure 5.**
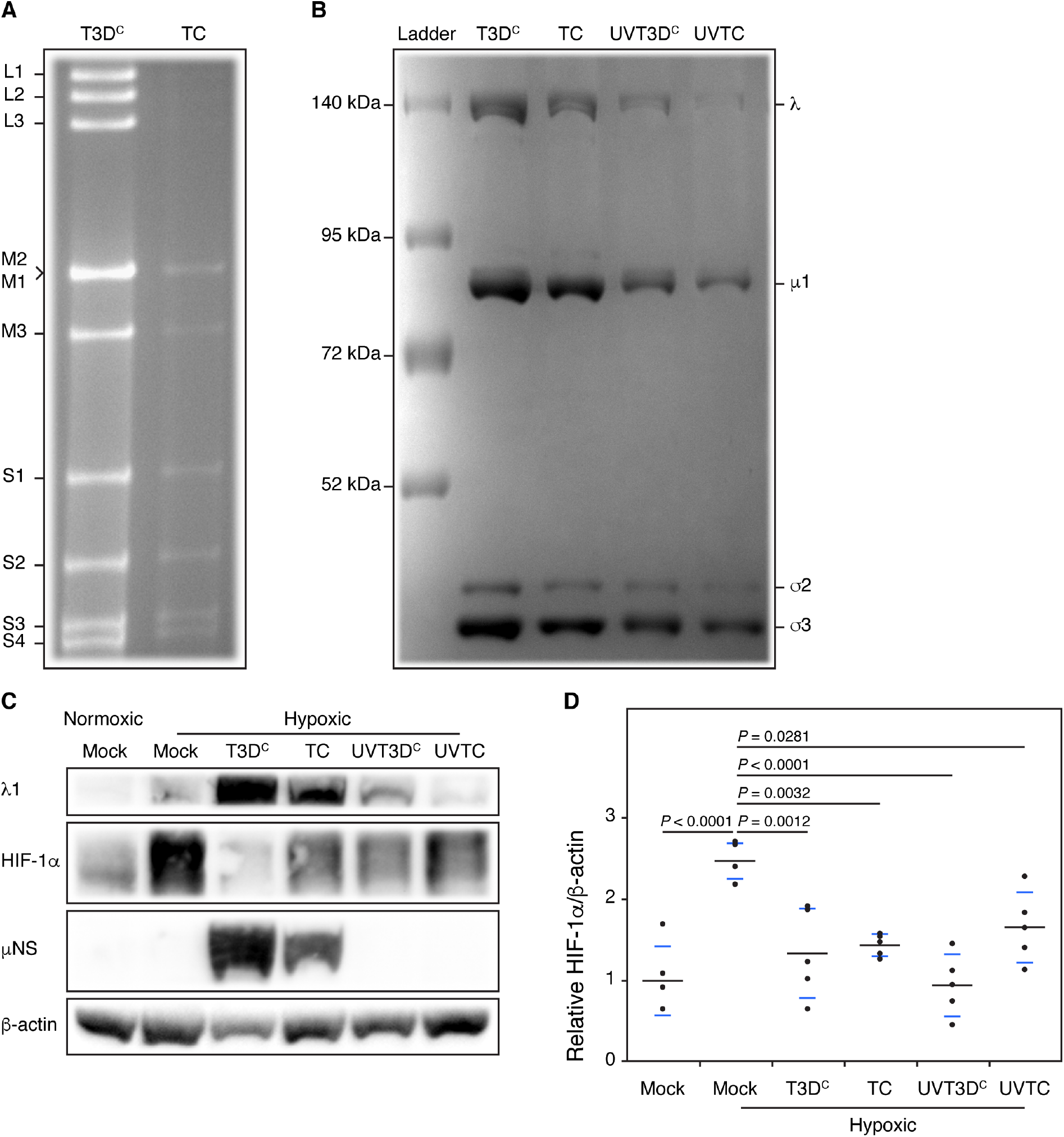
MRV dsRNA is not required for inhibition of HIF-1α accumulation. T3D^c^ virus was purified on a CsCl_2_ gradient and two bands were collected, a bottom highly infectious bottom component (T3D^c^) and a dsRNA deficient top component (TC). (A) 1.88 × 10^12^ particles of T3D^c^ and TC were subject to TRlzol RNA extraction and run on SOS-PAGE at 20 mA for 10 h to separate the 10 MRV genome segments. T30c and TC were UV-irradiated at 1 J/cm^2^ (UVT3D^c^ and UVTC), and (B) 1.88 × 10^12^ particles of each virus were run on SOS-PAGE followed by Coomassie staining to observe total structural proteins. PC3 cells were mock-infected or infected with T3D^c^, TC, UVT3D^c^, or UVTC, and at 20 h p.i. cells were exposed to normoxic or hypoxic conditions for 4 h and were collected for immunoblot analysis. (C) Cell lysates were run on SOS-PAGE, transferred to nitrocellulose and stained with antibodies against the virus structural protein λ1, the nonstructural μNS, HIF-1α and β-actin. (0) lmmunoblots were analyzed on Quantity One software to quantify HIF-1 α/β-actin protein levels. Black dots represent each replicate, the black line represents the average and the blue lines represent the standard deviation. *P* values were determined using ANOVA with Tukey’s multiple comparison test in JMP® Pro 14.2.0.

### UVT3D^C^-induced inhibition of HIF-1α is dependent on the proteasome

Throughout this work we observed significant differences between the impact of T3D^C^ and UVT3D^C^ on HIF-1α accumulation during infection, specifically measuring significant increases in the inhibition of HIF-1α accumulation in UVT3D^C^ infected cells. We reasoned that UV-inactivation may result in a virus that utilizes different mechanisms to inhibit HIF-1α compared to T3D^C^. Using an approach similar to that published for T3D^C^ (Gupta-Saraf and Miller, 2014), we investigated the role played by the proteasome in UVT3D^C^ inhibition of HIF-1α. Because prior work demonstrated that MRV inhibition of HIF-1α accumulation was proteasome-dependent at early times p.i., while translation inhibition was responsible for HIF-1α accumulation at late times p.i., we investigated this question at 12 h p.i. instead of the 24 h p.i. timepoint we used throughout the rest of this study. PC3 cells were infected with T3D^C^ or UVT3D^C^, and at 8 h p.i. cells were treated with the proteasome inhibitor, MG132, and placed under hypoxia for 4 h. At 12 h p.i. the cells were harvested and subjected to immunoblot analysis. In the absence of MG132, T3D^C^ and UVT3D^C^ inhibited HIF-1α as expected; however, in the presence of MG132, both viruses were prevented from inhibiting HIF-1α accumulation (Fig. 6A-B). This data suggests that despite the differences that we measured in efficiency of HIF-1α inhibition between T3D^C^ and UVT3D^C^, that like T3D^C^, UVT3D^C^ inhibits HIF-1α accumulation at least in part by inducing proteasome-mediated degradation of HIF-1α.

**Figure 6.**
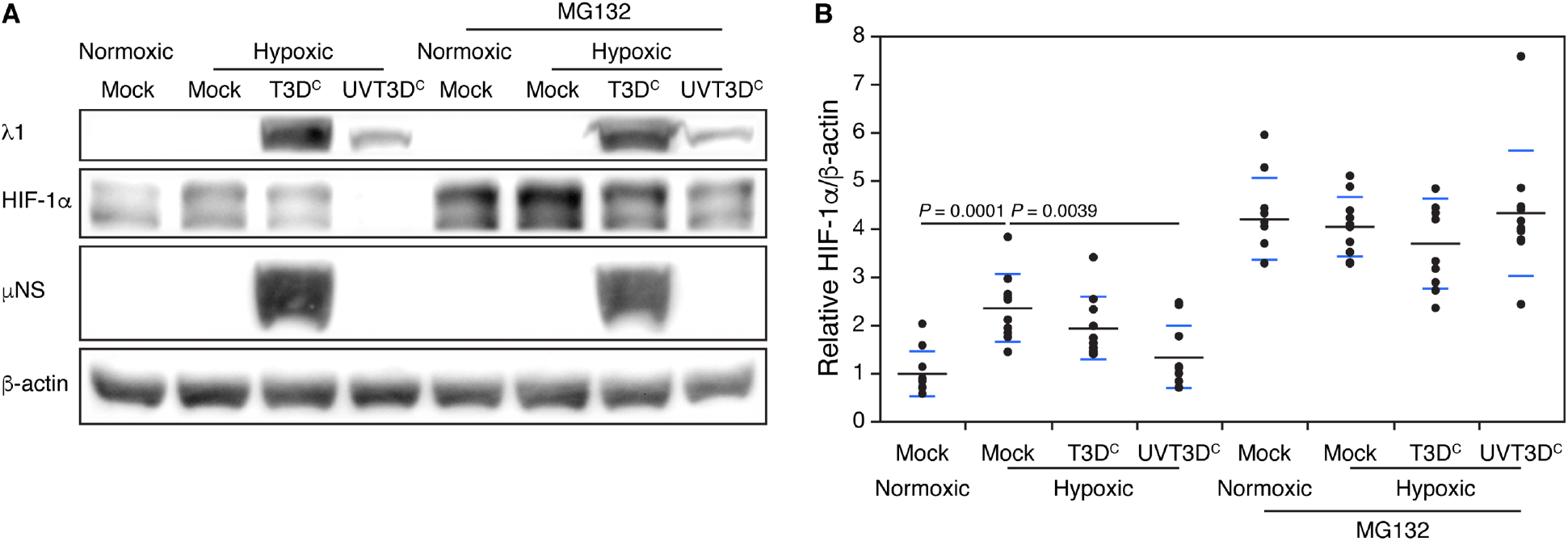
UVT3D^c^ inhibits HIF-1α through proteasome-mediated degradation early during infection. PC3 cells were mock-infected or infected at an MOI = 20 with T3D^c^ or an equal number of viral particles of T3D^c^ exposed to 1 J/cm^2^ of UV-irradiation (UVT3D^c^). At 8 h p.i. cells were treated with the proteasome inhibitor MG132 (50 μM) and exposed to normoxic or hypoxic conditions for 4 hat which point the cells were collected for immunoblot analysis. (A) Cell lysates were run on SOS-PAG E, transferred to nitrocellulose, and stained with antibodies against the virus structural protein λ1, the nonstructural μNS, HIF-1α and γ-actin. (B) lmmunoblots were analyzed on Quantity One software to quantify HIF-1α/β-actin protein levels. Black dots represent each replicate, the black line represents the average and the blue lines represent the standard deviation. *P* values were determined using ANOVA with Tukey’s multiple comparison test in JMP® Pro 14.2.0.

## DISCUSSION

Mammalian orthoreovirus inhibition of HIF-1α is an intriguing characteristic of the virus that could be exploited to help inhibit the devastating effects of hypoxia and HIF-1α accumulation in tumors. In this work, the stages of viral infection were dissected to better understand when MRV inhibits HIF-1α during its lifecycle. Experiments with NH_4_Cl, E64, and UV-inactivated virus suggest a step between viral capsid cleavage and transcription is sufficient for this phenotype. This step is concurrent with endosomal membrane disruption and release of the viral core and other endosomal contents, including cleaved σ3 and μ1, as well as a smaller amount of σ1, into the cytoplasm. (Dryden et al., 1993). Additionally, it has been proposed that UV-inactivation may destabilize the viral core and result in increased dsRNA release (Sherry, 2009). Therefore, in our work we focused on these viral components. It is evident that loss of the σ3 protein, μ1-induced apoptosis, or dsRNA, which has been suggested to play a role in HIF-1α inhibition in cells following transfection (Hotani et al., 2015), did not significantly impact T3D^C^-induced inhibition of HIF-1α during the natural route of infection. Taken together, this data suggests that an as yet unidentified consequence of capsid protein cleavage and/or endosomal membrane disruption is primarily responsible for inhibition of HIF-1α protein accumulation, independent of viral outer capsid proteins, σ3, the apoptosis-inducing function of μ1, or the presence of dsRNA.

While there is a possibility that the MRV cellular attachment protein, σ1, may inhibit HIF-1α, we find this possibility to be unlikely since there are only a maximum of 36 copies per virion, compared to 600 copies of the capsid proteins σ3 and μ1 (Dietrich et al., 2018). σ1 binds JAM-A and/or sialic acid to enter cells (Barton et al., 2001; Paul et al., 1989; van den Wollenberg et al., 2012), and following removal of σ3, and subsequent μ1 cleavage, the MRV capsid changes confirmation to release σ1 into the endosome. While it is possible that σ1-receptor binding could result in a downstream signal that inhibits HIF-1α, σ1-receptor engagement would still occur, and downstream signals would presumably still be present in experiments where endosomal protease inhibitors are present. In infected cells treated with NH_4_Cl or E64 we observed rescue of HIF-1α accumulation, suggesting any signaling that occurs as a result of σ1-receptor binding does not inhibit HIF-1α. It is also possible that release of the viral core into the cytoplasm may be sufficient to inhibit HIF-1α. Future work will investigate the extent by which the core and σ1 contribute to HIF-1α inhibition.

While we have not formally ruled out the possibility that σ1 or MRV core release is involved in MRV-induced HIF-1α inhibition, we find it more likely that the disruption of the endosome may contribute to HIF-1α inhibition. It is interesting that the ssDNA, non-enveloped human parvovirus (H-1 parvovirus), which is also endocytosed and requires low pH for escape from the late endosome, has been shown to inhibit HIF-1α accumulation (Cho et al., 2015). When H-1 parvovirus is treated with NH_4_Cl, HIF-1α is rescued from viral inhibition similar to what we have observed with MRV (Quattrocchi et al., 2012). The similarities between the requirements for HIF-1α inhibition between these two viruses may suggest that disruption of the endosome and subsequent downstream signaling may result in HIF-1α inhibition. Additionally, disruption of the late endosome, especially to the extent needed to release the large MRV core particle, likely results in release of endosomal contents into the cytoplasm, and while we focused on the viral components, the vesicles contain many other proteins and enzymes (Scott et al., 2014). These enzymes are not designed to be in the cytoplasm, and we reason that this release could result in various stress signals leading to inhibition of HIF-1α accumulation. In addition, late endosomes and lysosomes are targets for misfolded protein degradation, and the release of increased amounts of misfolded proteins may result in PERK activation which has previously been shown to result in HIF-1α inhibition (Ivanova et al., 2018; Jackson and Hewitt, 2016).

PERK is a part of the stress response of the cell and is responsible for eIF2α phosphorylation resulting in stress granule (SG) formation (Ron and Walter, 2007). SGs are aggregates of stalled mRNA-ribosome complexes, RNA-binding proteins, and SG effector proteins that form when cells experience stress (Anderson and Kedersha, 2009). It is interesting that UV-inactivated T3D^C^ inhibits HIF-1α significantly more than T3D^C^, because UV-inactivated virus also induces substantially higher numbers of SGs that are maintained for a prolonged period of time compared to infectious virus (Qin et al., 2009). MRV-induction of SGs is dependent on eIF2α phosphorylation, and correlates with inhibition of cellular, but not viral translation (Qin et al., 2011; Qin et al., 2009). Therefore, it is possible that the observed increase in HIF-1α inhibition induced by UVT3D^C^ may occur, at least in part, as a result of increased translational shutoff. It has also been shown that SGs can directly impact HIF-1α accumulation. As mentioned previously PERK activated SGs can inhibit HIF-1α, and under extreme hypoxia, the SG nucleating proteins TIAR and TIA-1 can form SGs that also suppress HIF-1α expression (Gottschald et al., 2010). Therefore, it is conceivable that endosomal disruption and release of the outer capsid proteins, cellular proteins, and core particle increases SG formation and stress signaling, leading to decreased accumulation of HIF-1α. Further research investigating the role of SGs in HIF-1α inhibition is needed.

In addition to analyzing the stage of viral replication sufficient to inhibit HIF-1α, our data also demonstrated that UVT3D^C^ induced inhibition of HIF-1α is dependent on proteasome-mediated degradation pathway early during infection, as seen in T3D^C^ infected cells (Gupta-Saraf and Miller, 2014). This suggests that UVT3D^C^ may employ the same mechanisms of HIF-1α inhibition as T3D^C^, but to a greater extent. This is important as it suggests that UVT3D^C^ may be a better candidate for inhibiting HIF-1α in hypoxic tumors. While research has shown that MRV inhibits HIF-1α activity *in vivo*, the UV-inactivated virus was unable to inhibit HIF-1α *in vivo* under the same conditions (Hotani et al., 2019). It is curious that UV-inactivation works significantly better *in vitro* but does not show any inhibition *in vivo*. The fact that UV-inactivated virus cannot replicate and infect new cells after initial infection like MRV likely explains why no inhibition was observed. In order for UV-inactivated virus to be useful in a clinical setting much higher titers of the virus would likely be necessary. While MRV may be useful as a treatment in many cancer patients, it is important to consider that active viral therapy may not be amenable to all patients. Therefore, our work has provided early evidence for further investigation into the use of MRV and UV-inactivated MRV in tumors to mitigate the detrimental effects associated with tumor hypoxia.

## ACKNOWLEDGMENTS

We thank Dr. Pranav Danthi for generously providing the T3D^F^ M2 (I595K) mutant. This work was supported by the Bailey Research Career Development Award and NIH R15 grant to C.L.M.

